# MRI-based 3D Estimation of Skeletal Muscle Architecture and Strain during Contraction

**DOI:** 10.1101/2025.07.23.666431

**Authors:** Roberto A. Pineda Guzman, Carly A. Lockard, Xingyu Zhou, Evelyn Bombard, Crystal Coolbaugh, Mariana E. Kersh, Bruce M. Damon, Melissa T. Hooijmans

**Author notes:** Address for Correspondence: Bruce M. Damon, PhD, Stephens Family Clinical Research Institute, Carle Foundation Hospital, 611 W. Park Ave., Urbana, IL 61801.

## Abstract

Skeletal muscle generates forces that drive the motion of the human body. Three-dimensional (3D) quantification of whole-muscle architecture and strain, and their relationship during contraction is critical to understanding the mechanical function of skeletal muscle in health and disease. This has proven to be challenging, as brightness mode ultrasound is capable of measuring muscle architecture during contraction but cannot capture 3D changes in whole-muscle architecture, while Diffusion Tensor Imaging (DTI)-based tractography can measure 3D whole-muscle architecture but its use during contraction is precluded by long scan times (>5 minutes). In this study, we implement DTI-based tractography with an image registration-based approach, previously validated under passive deformation, to estimate 3D whole-muscle architecture of the tibialis anterior (TA) muscle during moderate intensity contractions (20-40% MVC). Moreover, this approach allows the measurement of whole-muscle strain during contraction, facilitating the evaluation of intramuscular relationships between architecture and strain. Our results show a decrease in the fiber-tract length, an increase in the pennation angle, and an increase in the fiber curvature of the TA during contraction. Intramuscular strain heterogeneity was observed between and within different regions of the muscle, with exploratory analyses suggesting that regional strain heterogeneity could be influenced by muscle architecture. Our results showcase the potential of MRI-based methods to obtain 3D estimates of whole-muscle architecture and strain during contraction, providing a breadth of new data that allows for new avenues of skeletal muscle biomechanical research.

## 1. Introduction

Skeletal muscle generates forces that drive the motion of the human body. Muscle architecture – the internal arrangement of the muscle’s fibers with respect to its line of action – is a key structural determinant of this function (Lieber & Friden, 2000). Muscle fiber length and orientation influence a muscle’s force production, excursion (Lieber & Friden, 2000), and lengthening or shortening velocity (Sacks & Roy, 1982). Moreover, muscle fiber orientation and curvature affect intramuscular strain development during contraction (Azizi & Deslauriers, 2014; Blemker et al., 2005), affecting the muscle’s injury susceptibility (Silder et al., 2010). Thus, three-dimensional (3D) quantification of whole-muscle architecture, strain, and their relationship during contraction is key to understanding the structural mechanisms underlying the mechanical function of skeletal muscle in health and disease.

Brightness-mode ultrasound is the most widely used method to measure local muscle architecture during contraction (Ito et al., 1998; Raiteri et al., 2016), but cannot capture 3D changes in whole-muscle architecture (Van Hooren et al., 2020), leaving an unmet need for methods capable of estimating 3D whole-muscle architecture during contraction. Alternatively, 3D measurements of whole-muscle architecture can be obtained from Diffusion Tensor Imaging (DTI), a magnetic resonance imaging (MRI) method that exploits the correspondence between the local muscle fiber orientation and the direction of greatest diffusion (Cleveland et al., 1976), as represented by the first eigenvector of the diffusion tensor (Van Donkelaar et al., 1999). The first eigenvector is integrated to create “fiber-tracts” that represent muscle architecture at the spatial scale of several fascicles (Damon et al., 2017). Skeletal muscle architecture can be estimated from DTI-based tractography at rest (Bolsterlee et al., 2018; Damon et al., 2002; Froeling et al., 2012), providing quantitative estimates of muscle fiber length (Heemskerk et al., 2005), pennation angle (Lansdown et al., 2007), and fiber curvature (Damon et al., 2012). However, long scan times (> 5 minutes) preclude the use of DTI during contraction.

A promising alternative to measure muscle architecture during contraction is DTI-based tractography coupled with an image registration approach. (Hooijmans et al., 2025) found that displacement fields generated from the registration of MRI structural images can be used to transform the DTI-based fiber-tracts from an undeformed to a passively deformed state, with mean architecture parameters that do not differ between the transformed fiber-tracts and those measured in the deformed state. This approach may be adapted to the estimation of muscle architecture during contraction. Moreover, image registration has been previously used to measure muscle strain during passive deformation (Pamuk et al., 2016; Yaman et al., 2013) and contraction (Karakuzu et al., 2017). Structural images with whole-muscle coverage can be acquired in <40 s, allowing for whole-muscle deformation data to be obtained from a single repetition of a moderate intensity contraction, defined here as 20-40% of the maximum voluntary contraction (MVC) force. This feature is unavailable in other commonly used deformation-mapping techniques such as spatial-tagging (Englund et al., 2011) and phase-contrast MRI (Hooijmans et al., 2024; Malis et al., 2020; Mazzoli et al., 2018), which have the temporal resolution to characterize contraction development over time but require multiple contraction repetitions to acquire whole-muscle data.

The objective of this study was to implement a registration-based approach of DTI and structural images to quantify whole-muscle architecture and strain during isometric contraction. Focusing on the tibialis anterior muscle, we hypothesized that i) muscle fiber-tract length would decrease and pennation angle and fiber curvature would increase during isometric contraction; and ii) strain would develop heterogeneously along and within different muscle regions. Additionally, we explored correlations between muscle architecture and strain development during contraction.

## 2. Methods

### 2.1 Experimental protocol and image acquisition

Data collection for this IRB-approved study occurred at Vanderbilt University Medical Center. The data were transferred to Carle Foundation Hospital under a Data Use Agreement, and data analysis continued under local IRB approval. Seven healthy participants (5 male, 2 female, age range: 23 – 31 years) provided written informed consent. All participants received an orientation session with an MRI compatible isometric dorsiflexion device that measures dorsiflexion forces (Maguire et al., 2007). The participants performed maximal isometric dorsiflexion, and the MVC force was measured as the mean force during three contractions having a <5% difference in their maximal force. Two submaximal contraction forces at 20% and 40% of the MVC force were determined for the MRI session.

During the MRI session, the participants laid feet-first supine in a 3 tesla MR system (Philips Achieva dStream, the Netherlands), with their right ankle and foot secured in the dorsiflexion device at 10° of plantarflexion. MRI datasets of the right lower leg were acquired using a 16-element receiver coil and the 12-element receiver coil built into the patient table. The 16-element coil was placed on top of the lower legs with foam pillows and fixation bands to avoid muscle compression and coil movement during contractions.

DTI data were acquired to estimate muscle fiber direction maps and muscle architecture during rest. The DTI data acquisition included two separate image stacks, each using a diffusion weighted spin-echo echo-planar imaging (SE-EPI) sequence (Table 1). The two image stacks had an overlap of four slices (28 mm) and spanned a foot-head distance of 308 mm, covering the lower leg from the ankle to the tibial plateau.

**Table 1.**
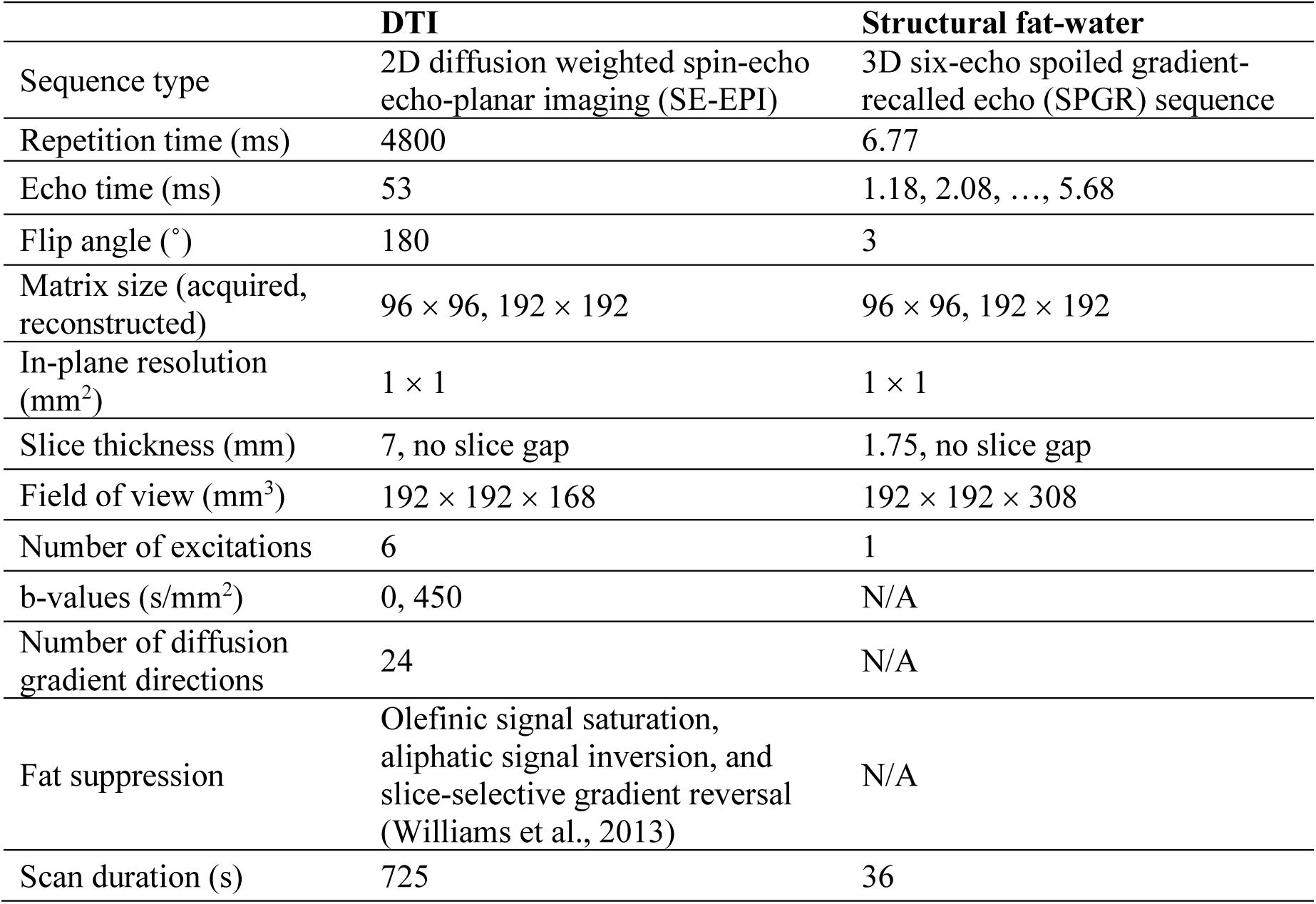
MRI sequence parameters used in this study.

Fat-water images covering the combined field of view of the DTI stacks were obtained for anatomical reference and image registration. The images were acquired using a 3D six-echo spoiled gradient-recalled echo (SPGR) sequence (Table 1) with a scan duration of 36 seconds. After acquiring images at rest, the participants were asked to hold dorsiflexion contractions for 45 seconds at 20% and 40% MVC with the help of real-time visual feedback to match the target force. While the participants held the contraction, an image of the lower leg was acquired. A second set of images were then acquired at rest and during contraction. The order of the contraction intensities (20% and 40% MVC) was randomized for each participant.

### 2.2 Image data analysis

Figure 1 provides an overview of the image data analysis workflow. All analyses were performed in MATLAB R2023b using the publicly available *MuscleDTI_Toolbox* (Damon et al., 2021) and custom scripts.

**Figure 1.**
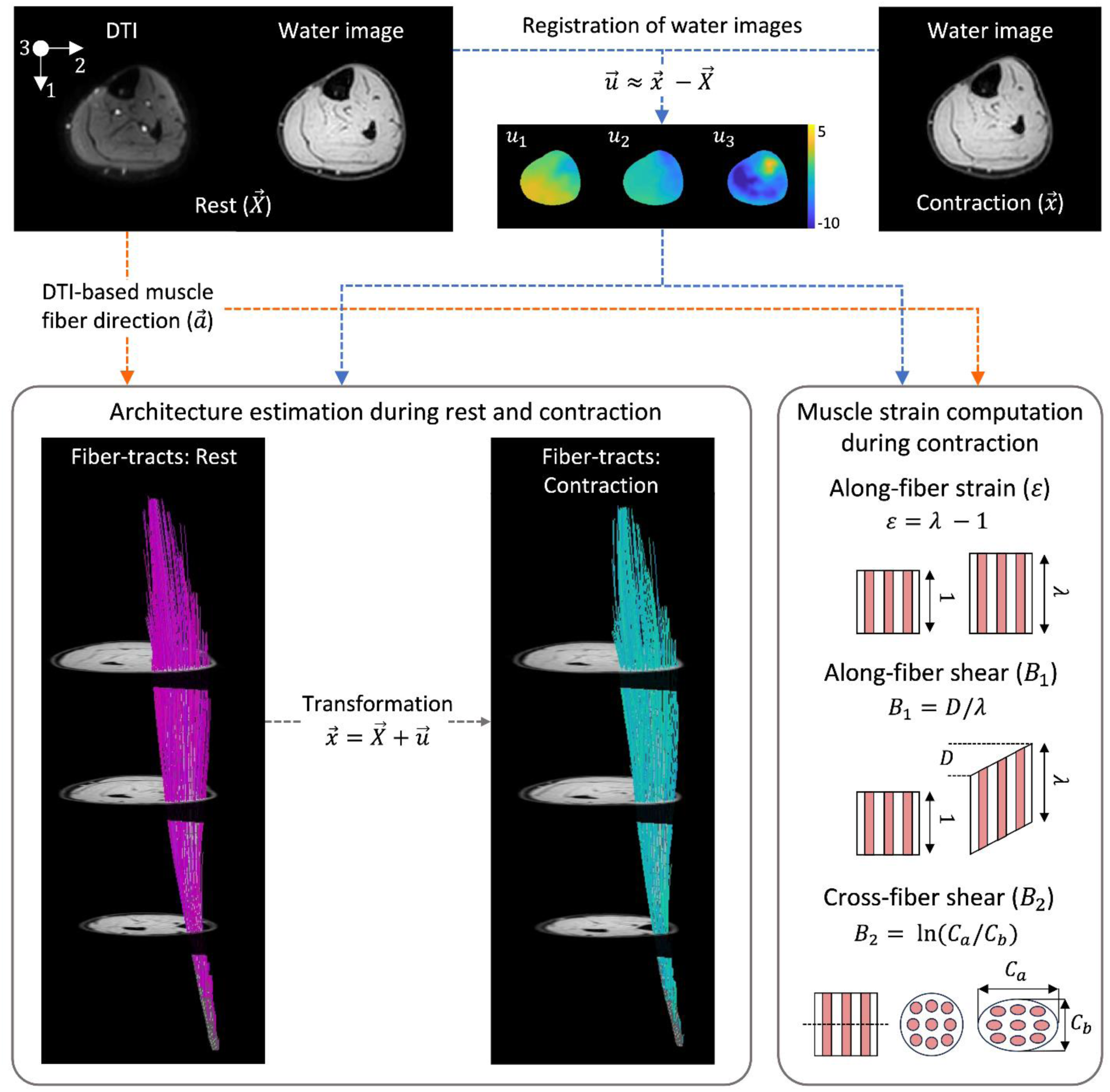
Overview of the image data analyses conducted in the study. DTI data was acquired during rest while fat-water structural images were acquired during rest prior to the contractions and during contraction at 20% and 40% MVC, where anterior bulging of the TA can be observed. Registration of the water images was used to estimate the displacement field 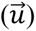 mapping the lower leg from rest (undeformed configuration, 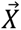) to contraction (deformed configuration, 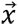). The three-dimensional components of 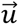 are shown, with units in millimeters. The muscle fiber direction field 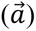 was obtained from the DTI data during rest. Fiber-tracts representing muscle architecture were computed during rest from 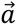, and were transformed to their contracted (deformed) configuration with 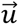. Muscle strain invariants during contraction were computed from 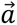 and 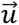. Along-fiber strain (*ε*) is a measure of the change in length from one to *λ* in the representative muscle unit element, along-fiber shear (*B*_1_) is a measure of the change in shape parallel to the muscle fibers by a distance D across the horizontal axis of the unit element, and cross-fiber shear (*B*_2_) represents the change in ellipticity of a circular muscle element with unit diameter.

#### 2.2.1 Estimation of muscle fiber direction and architecture during rest

A single DTI 44-slice dataset was obtained after concatenating the two DTI image stacks and eliminating the redundant copies of the overlapping slices. Each slice of the DTI dataset was registered to its corresponding slice of the structural rest images using 2D Demons-based nonrigid registration (Lockard et al., 2024). The registered DTI dataset was denoised with anisotropic smoothing at a noise level of 5% (Buck et al., 2015; Ding et al., 2005; Xu et al., 2010). A weighted linear-least squares algorithm was used to compute the diffusion tensor in each voxel of the denoised images. The diffusion tensor was diagonalized using singular value decomposition. The muscle fiber direction field 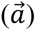 was computed from the first eigenvector of the diffusion tensor, while fractional anisotropy (FA) was computed from the tensor’s eigenvalues. The 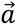 was smoothed using a penalized least squares approach combined with the discrete cosine transform (Garcia, 2010, 2011, 2020), using a smoothing parameter of 5.

The tibialis anterior (TA) muscle and its compartments were manually segmented from the water images (Supplementary Figure 1) and used in all further analyses. DTI fiber-tracts of the TA were generated using an aponeurosis-based seeding method (Damon et al., 2024), with the seed-points defined in a 30×20 mesh created from the segmented muscle aponeurosis. Fiber-tracts were generated using the Euler integration of 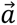 with a step size of 1 mm. Tracts were terminated at the boundary of the muscle mask, at points where FA fell outside a range of 0.05-0.40, or when the angle between consecutive points exceeded 30°. Fiber-tracts were smoothed using 3^rd^ order polynomial fitting (Lockard et al., 2024). Fiber-tract length (L_FT_), pennation angle (α), and curvature (κ) were computed for each fiber-tract (Damon et al., 2021; Damon et al., 2012; Lansdown et al., 2007). Fiber-tracts shorter than 10 mm were omitted. The whole-muscle mean architecture parameters were computed for L_FT_, α, and κ. The fiber-tracts corresponding to the *Deep* and *Superficial* TA compartments were generated using manually segmented masks from each compartment and their mean architecture parameters were computed. To represent the architectural variation within each muscle, fiber-tracts were grouped into fiber-tract clusters with structurally similar fiber-tracts using the algorithm presented in (Damon et al., 2002), and the mean architecture parameters of each cluster were computed. The Supplementary materials provide details of the fiber-tract clustering method.

#### 2.2.2 Estimation of muscle deformation during contraction

The displacement field 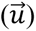 mapping the rest (undeformed configuration, 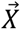) to the contraction (deformed configuration, 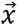) images was estimated via additive 3D Demons-based registration using MATLAB’s *imregdemons* function (Thirion, 1998; Vercauteren et al., 2009). Water images derived from the SPGR acquisitions were registered using an *AccumulatedFieldSmoothing* of 1.5 and 4 *PyramidLevels*. A detailed description of the determination of the image registration parameters is available in the Supplementary material. The displacement field was then smoothed using a 3×3×3 Gaussian kernel over 10 iterations to improve the precision of strain estimates (Chan et al., 2013) and resized to match the resolution of the DTI data.

#### 2.2.3 Estimation of muscle architecture during contraction

Muscle fiber-tracts were transformed to their contracted state 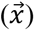 by adding 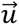 to the position vectors describing the fiber-tracts and aponeurosis mesh during rest 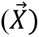 (Hooijmans et al., 2025). The transformed fiber-tracts longer than 10 mm were preserved and smoothed using 3^rd^ order polynomial fitting, and mean L_FT_, α, and κ were computed for the whole TA muscle and its *Deep* and *Superficial* compartments.

#### 2.2.4 Muscle strain computations

Physically based strain metrics (Blemker et al., 2005; Criscione et al., 2001) of the TA during contraction were computed from 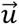. The deformation gradient ***F*** and the right Cauchy-Green strain tensor ***C*** were computed in each voxel with

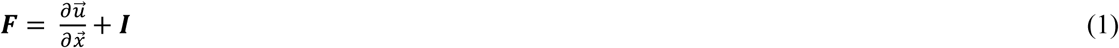

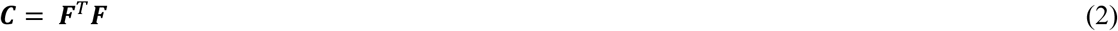

where ***I*** is the 2^nd^ order identity tensor. The invariants of ***C*** accounting for transverse isotropy perpendicular to the muscle fiber direction, 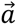, were computed as

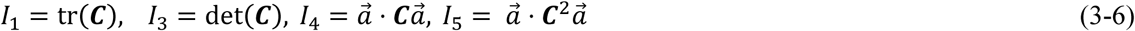

Along-fiber strain (*ε*), along-fiber shear (*B*_1_), and cross-fiber shear (*B*_2_) were computed in each voxel and in each point of the smoothed fiber-tracts using the equations listed in Table 2. The physical interpretation of each strain metric is sketched in Figure 1. In the fiber-tract points, muscle fiber direction 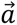 was estimated from the tangent vector of each point. The mean strain metrics were computed in three distinct regions of the muscle, two regions corresponding to the *Deep* and *Superficial* compartments, and the *No-Aponeurosis* region defined as the portion of the muscle lying superior to the aponeurosis (Supplementary figure 1). The *No-Aponeurosis* region was defined to characterize the distinct strain patterns observed in this muscle region during preliminary analyses. The strain metrics were also analyzed longitudinally by computing the mean invariants of each slice and normalizing the measurements to the region’s length. For the fiber-tract analyses, the mean strain metrics were computed in each fiber-tract cluster by averaging the mean strain metrics of all the fiber-tracts.

**Table 2.**
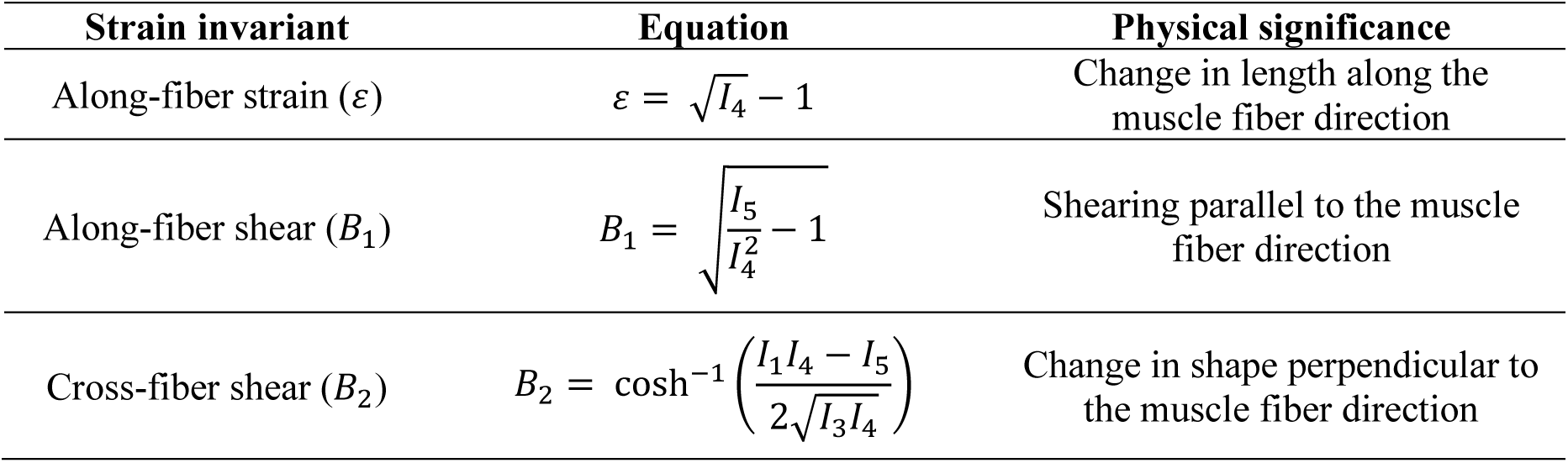
Physically based strain invariants measured in the muscle during contraction.

### 2.3 Statistical analysis

Statistical analyses were conducted in R (version 4.3.3). Data normality was verified using a Shapiro-Wilks test for architecture parameters and quantile-quantile plots for the mean compartment strain metrics. The differences between the architecture parameters of the whole muscle and each compartment during rest and contraction at 20% and 40% MVC were evaluated using paired t-tests with Holm-Bonferroni correction for multiple comparisons. Two-way Analysis of Variance (ANOVA) was conducted to evaluate the effect of muscle region and contraction intensity in the muscle strain metrics, accounting for the participants as a random effect. Post-hoc comparisons were conducted using estimated marginal means with a Holm-Bonferroni correction. Associations between the strain metrics and architecture parameters of the fiber-tract clusters were exploratively evaluated using repeated measures correlations (Bakdash & Marusich, 2017), considering the different fiber-tract clusters of each participant as repeated measures. The significance level was set at 0.05.

## 3. Results

The mean ± standard deviation of the MVC forces generated by the participants during dorsiflexion was 265.3 ± 130.1 N. In the 20% MVC target contractions, 19.1 ± 2.6% of the MVC force was generated by the participants, while 35.4 ± 0.6% of the MVC was generated in the 40% MVC target contractions.

### 3.1 Muscle architecture during contraction

Summary statistics of the architecture parameters measured during rest and contraction at 20% and 40% MVC are listed in Table 3 and shown in Figure 2. Mean whole-muscle fiber-tract length (L_FT_) decreased by 6% (*p*<0.001) during 20% MVC, and by 12% (*p*<0.001) during 40% MVC. L_FT_ decreased by 6% in the *Deep* compartment (*p*=0.002) and 5% in the *Superficial* compartment (*p*=0.003) during 20% MVC and by 10% in the *Deep* compartment (*p*=0.001) and 6% in the *Superficial* compartment (*p*=0.003) during 40% MVC. The mean whole-muscle pennation angle (α) increased by 8% during 20% MVC (*p*<0.001) and 11% during 40% MVC (*p*=0.015). A 6% increase in α occurred in the *Deep* (*p*<0.001) and *Superficial* (*p*=0.042) compartments during 20% MVC, while a 10% increase in the *Deep* compartment (*p*=0.015) occurred at 40% MVC. Mean whole-muscle curvature (κ) increased by 39% (*p*=0.003) and 48% (*p*<0.001) during 20% and 40% MVC, respectively. In the Deep compartment, κ increased by 43% (*p*=0.004) and 61% (*p*=0.001) during 20% and 40% MVC, compared to increases of 27% (*p*=0.019) and 33% (*p*=0.013), respectively, in the *Superficial* compartment.

**Figure 2.**
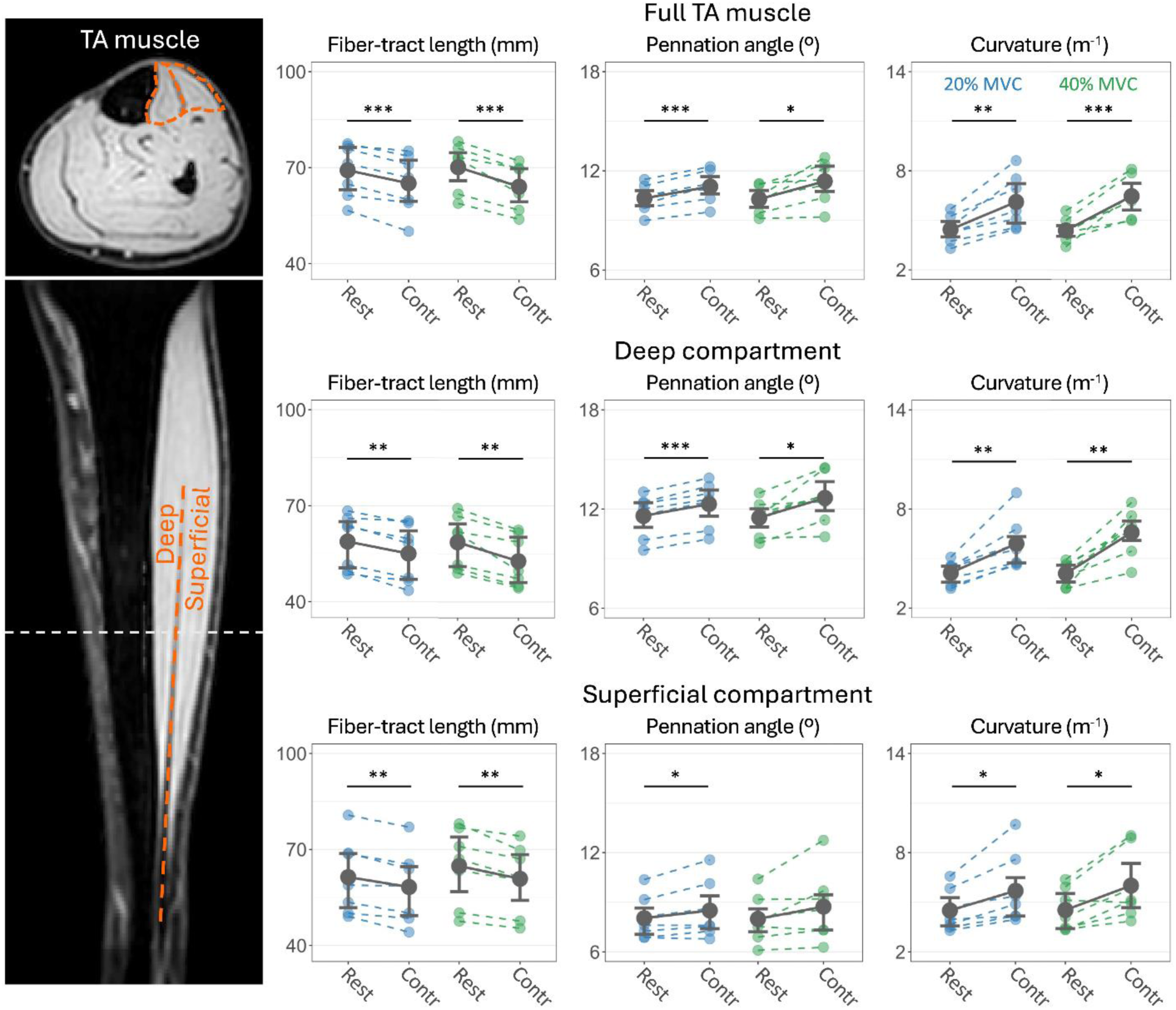
Mean architecture parameters of the tibialis anterior muscle measured during rest and contraction at 20% and 40% MVC. Each dot represents the data from each participant, while the lines connect the measurements obtained from the same participant. Data for the full muscle is shown in the top row, for the *Deep* compartment in the middle row, and for the *Superficial* compartment in the bottom row. A decrease in fiber length and an increase in pennation angle and curvature was observed during 20% MVC (blue) and 40% MVC (green). The water image on the left shows a longitudinal view of the TA muscle and the compartmental division of the muscle that divided the fiber-tracts based on their origin in the aponeurosis (dashed line). \**p* < 0.05, \*\**p* < 0.01, \*\*\**p* < 0.001

**Table 3.** Summary statistics (mean ± standard deviation) of the architecture parameters measured in the rest and contracted state for the 20% and 40% MVC contractions.

| Architecture parameter |  | Rest | 20% MVC | *p* | Rest | 40% MVC | *p* |
| --- | --- | --- | --- | --- | --- | --- | --- |
| Fiber-tract length, $L_{FT}$ (mm) | Full TA | 69.2 ± 8.3 | 65.1 ± 9.2 | **<0.001** | 70.1 ± 7.3 | 64.0 ± 7.0 | **<0.001** |
| | Deep | 58.8 ± 8.5 | 55.0 ± 9.0 | **0.002** | 58.6 ± 8.2 | 52.7 ± 8.0 | **0.001** |
| | Superficial | 61.4 ± 11.8 | 58.3 ± 11.5 | **0.003** | 64.8 ± 12.1 | 60.8 ± 10.9 | **0.003** |
| Pennation angle, $\alpha$ ($^{\circ}$) | Full TA | 10.3 ± 0.8 | 11.1 ± 1.0 | **<0.001** | 10.3 ± 0.8 | 11.4 ± 1.3 | **0.015** |
| | Deep | 11.6 ± 1.3 | 12.3 ± 1.4 | **<0.001** | 11.5 ± 1.1 | 12.7 ± 1.5 | **0.015** |
| | Superficial | 8.0 ± 1.3 | 8.5 ± 1.8 | **0.042** | 8.0 ± 1.4 | 8.7 ± 2.1 | 0.092 |
| Curvature, $\kappa$ ($m^{-1}$) | Full TA | 4.4 ± 0.8 | 6.1 ± 1.6 | **0.003** | 4.4 ± 0.7 | 6.5 ± 1.2 | **<0.001** |
| | Deep | 4.1 ± 0.7 | 5.9 ± 1.6 | **0.004** | 4.1 ± 0.7 | 6.6 ± 1.4 | **0.001** |
| | Superficial | 4.5 ± 1.3 | 5.7 ± 2.2 | **0.019** | 4.5 ± 1.3 | 6.0 ± 2.1 | **0.013** |

### 3.2 Muscle strain metrics during contraction

Strain metrics measured during 20% and 40% MVC in each compartment are shown in Figure 3, with summary statistics provided in Table 4. There was a significant interaction between contraction intensity and muscle compartment in along-fiber strain, *ε* (*p*=0.001). *ε* ranged between −0.030 and 0.005 during 20% MVC, with no significant differences between the muscle regions. At 40% MVC, there was a higher *ε* in the *No-Aponeurosis* region than in the *Deep* (*p*=0.046) and *Superficial* (*p*=0.009) regions and a decrease in the *Superficial* region when compared to 20% MVC (*p*=0.036). Along-fiber shear (*B*_1_) was affected by contraction intensity (*p*=0.021), with a mean *B*_1_ across all regions of 0.110 at 20% MVC and 0.130 at 40% MVC. Moreover, there was a significant interaction between contraction intensity and muscle region (*p*=0.009) in *B*_1_, with a lower *B*_1_ in the *No-Aponeurosis* region than in the *Deep* region at 20% MVC (*p*=0.021). Cross-fiber shear (*B*_2_) was significantly affected by contraction intensity (*p*=0.001) and muscle region (*p*<0.001). *B*_2_ was higher in the *Deep* region than in the *Superficial* (*p*=0.021, *p*=0.026) and *No-Aponeurosis* (*p*=0.001, *p*=0.001) regions at 20% and 40% MVC, respectively. *B*_2_increased from 20% to 40% MVC in all three regions (*Deep* (*p*=0.021), *Superficial* (*p*=0.004), and *No-Aponeurosis* (*p*=0.021)).

**Figure 3.**
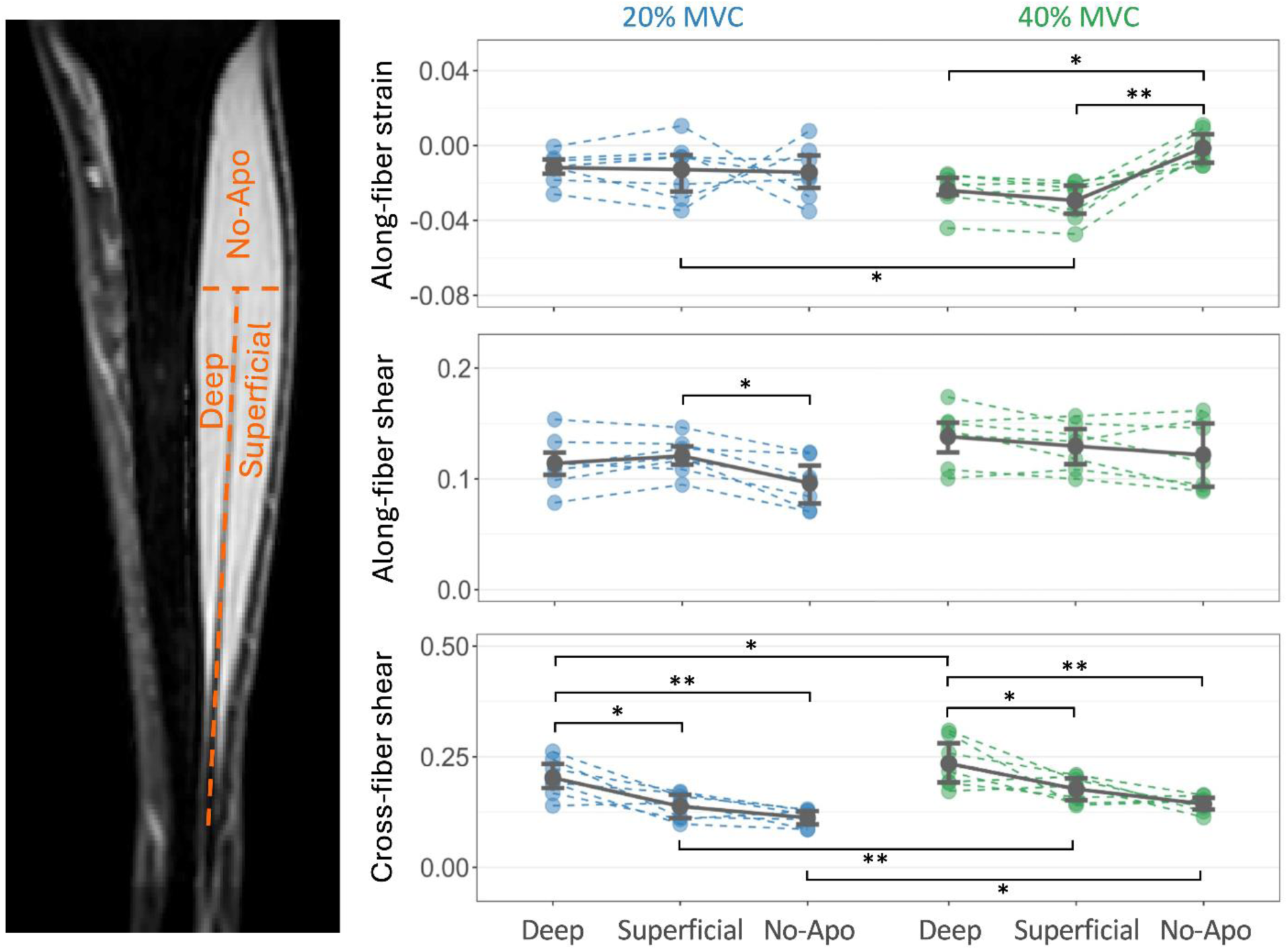
Mean along-fiber strain (top row) along-fiber shear (middle row) and cross-fiber shear (bottom row) measured in the different muscle regions at two different contraction intensities (20% and 40% MVC). Each dot represents the data from each participant, while the lines connect the measurements obtained from the same participant. Differences can be observed between the different regions in the same contraction intensity and between the two contraction intensities in the same region. The water image on the left shows how the muscle is divided into three different regions. The *No-Aponeurosis* region is referred to as No-Apo in this figure. \**p* < 0.05, \*\**p* < 0.01, \*\*\**p* < 0.001

**Table 4.** Summary statistics (mean ± standard deviation) of the strain metrics measured in the TA muscle during the 20% and 40% MVC contractions.

| Strain metrics |  |  |  | *p*-values (multiple comparisons) |  |  |  |
| --- | --- | --- | --- | --- | --- | --- | --- |
|  |  | 20% MVC | 40% MVC | 20% vs.<br>40% MVC |  | 20%<br>MVC | 40%<br>MVC |
| Along-fiber strain | Deep | -0.012 ± 0.009 | -0.024 ± 0.010 | 0.215 | Deep vs. Superficial | 1.000 | 1.000 |
| | Superficial | -0.013 ± 0.016 | -0.029 ± 0.011 | **0.036** | Superficial vs. No-Aponeurosis | 1.000 | **0.009** |
| | No-Aponeurosis | -0.014 ± 0.015 | -0.001 ± 0.009 | 0.166 | Deep vs. No-Aponeurosis | 1.000 | **0.046** |
| Along-fiber shear | Deep | 0.114 ± 0.024 | 0.138 ± 0.026 | 0.476 | Deep vs. Superficial | 1.000 | 1.000 |
| | Superficial | 0.121 ± 0.017 | 0.130 ± 0.021 | 1.000 | Superficial vs. No-Aponeurosis | **0.021** | 1.000 |
| | No-Aponeurosis | 0.096 ± 0.022 | 0.122 ± 0.032 | 0.433 | Deep vs. No-Aponeurosis | 0.160 | 0.214 |
| Cross-fiber shear | Deep | 0.203 ± 0.043 | 0.235 ± 0.056 | **0.021** | Deep vs. Superficial | **0.021** | **0.026** |
| | Superficial | 0.138 ± 0.031 | 0.177 ± 0.029 | **0.004** | Superficial vs. No-Aponeurosis | 0.480 | 0.287 |
| | No-Aponeurosis | 0.112 ± 0.019 | 0.143 ± 0.019 | **0.021** | Deep vs. No-Aponeurosis | **0.001** | **0.001** |

Longitudinal heterogeneity in the strain metrics was observed in all compartments (Figure 4). The mean *ε* of the seven participants was positive (tensile) in the inferior 20% length of the *Deep* and *Superficial* regions, and negative (compressive) in both regions in 50-80% length. In the *No-Aponeurosis* region, *ε* was compressive (indicating shortening) in the inferior 40% length and tensile (indicating lengthening) in the superior 50% length. *B*_1_and *B*_2_reached peak values at 25-65% length of the *Deep* and *Superficial* regions and decreased superiorly into the *No-aponeurosis* region.

**Figure 4.**
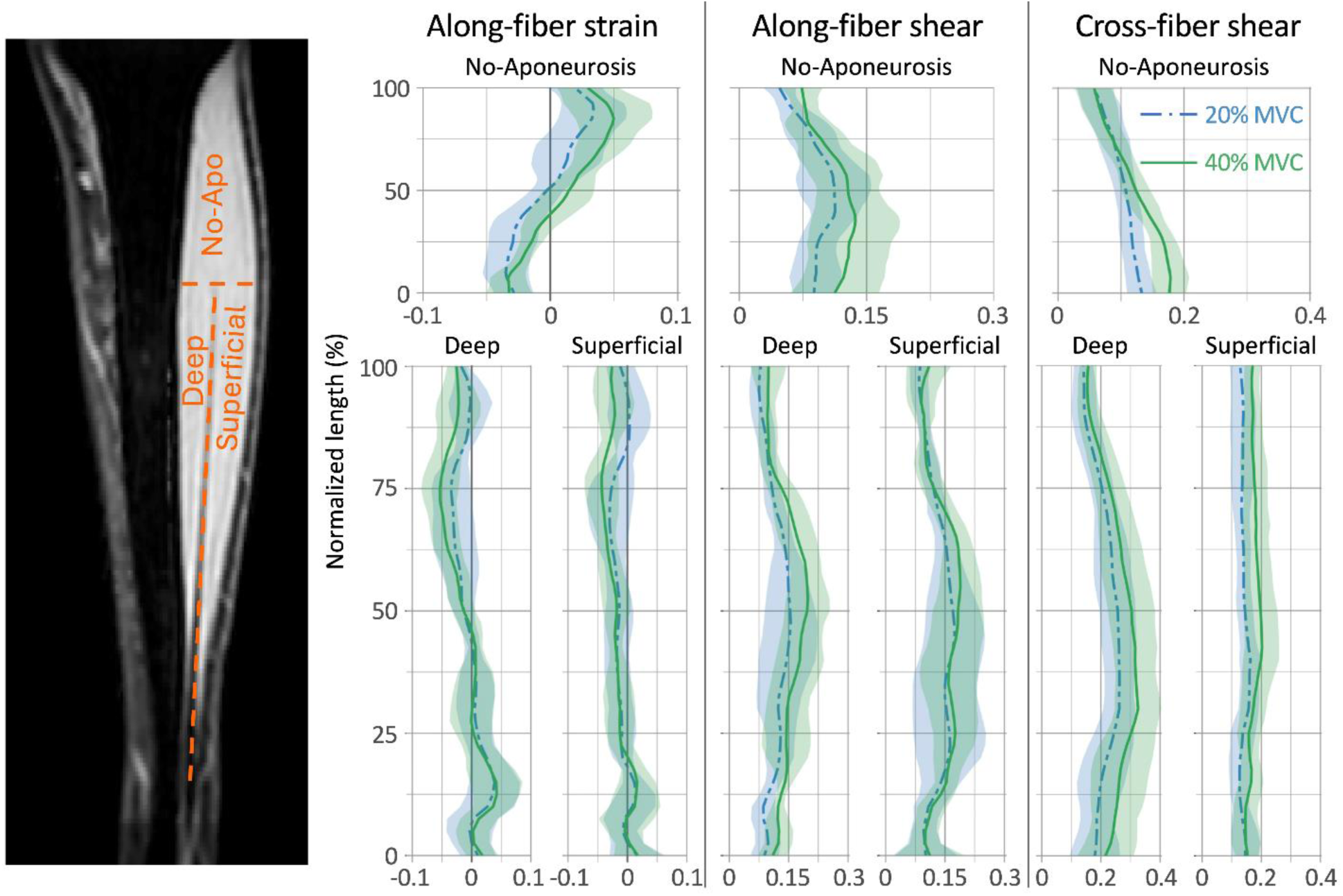
Mean along-fiber strain, along-fiber shear and cross-fiber shear of each axial slice of the muscle regions measured at two different contraction intensities (20% (blue) and 40% (green) MVC). The lines represent the mean strain for all participants in each slice, while the bounds represent the standard deviation. Tensile along-fiber strains (left panel) are observed in the inferior 25% of the *Deep* and *Superficial* regions, while compressive strains are observed in the superior 25% of the *Deep* and *Superficial* region and the inferior 50% of the *No-Aponeurosis* (No-Apo) region. The superior half of the *No-Aponeurosis* region shows tensile strains. Along-fiber shear reaches its peak around the 50% normalized length of the *Deep* and *Superficial* regions; and decreases until reaching the superior part of the muscle. Cross-fiber shear reaches its peak between the 25% and 50% normalized length of the *Deep* region, decreasing until reaching the superior part of the muscle.

### 3.3 Correlations between fiber-tract cluster architecture and strain metrics

TA muscles were divided into 3-5 fiber-tract clusters, depending on the architectural similarity within their fiber-tracts. Summary statistics of the repeated measures correlations are shown in Table 5. Mean fiber-tract cluster *ε* was significantly correlated with κ (*r*=0.40, *p*=0.045) during 20% MVC, and with α (*r*=-0.39, *p*=0.049) during 40% MVC. *B*_1_was significantly correlated with L_FT_ (*r*=0.40, *p*=0.045), α (*r*=0.55, *p*=0.004), and κ (*r*=0.71, *p*<0.001) during 20% MVC, and with L_FT_ (*r*=-0.48, *p*=0.015) and α (*r*=0.43, *p*=0.033) during 40% MVC. Plots depicting variable pairs with at least one significant correlation at 20% or 40% MVC are shown in Figure 5.

**Figure 5.**
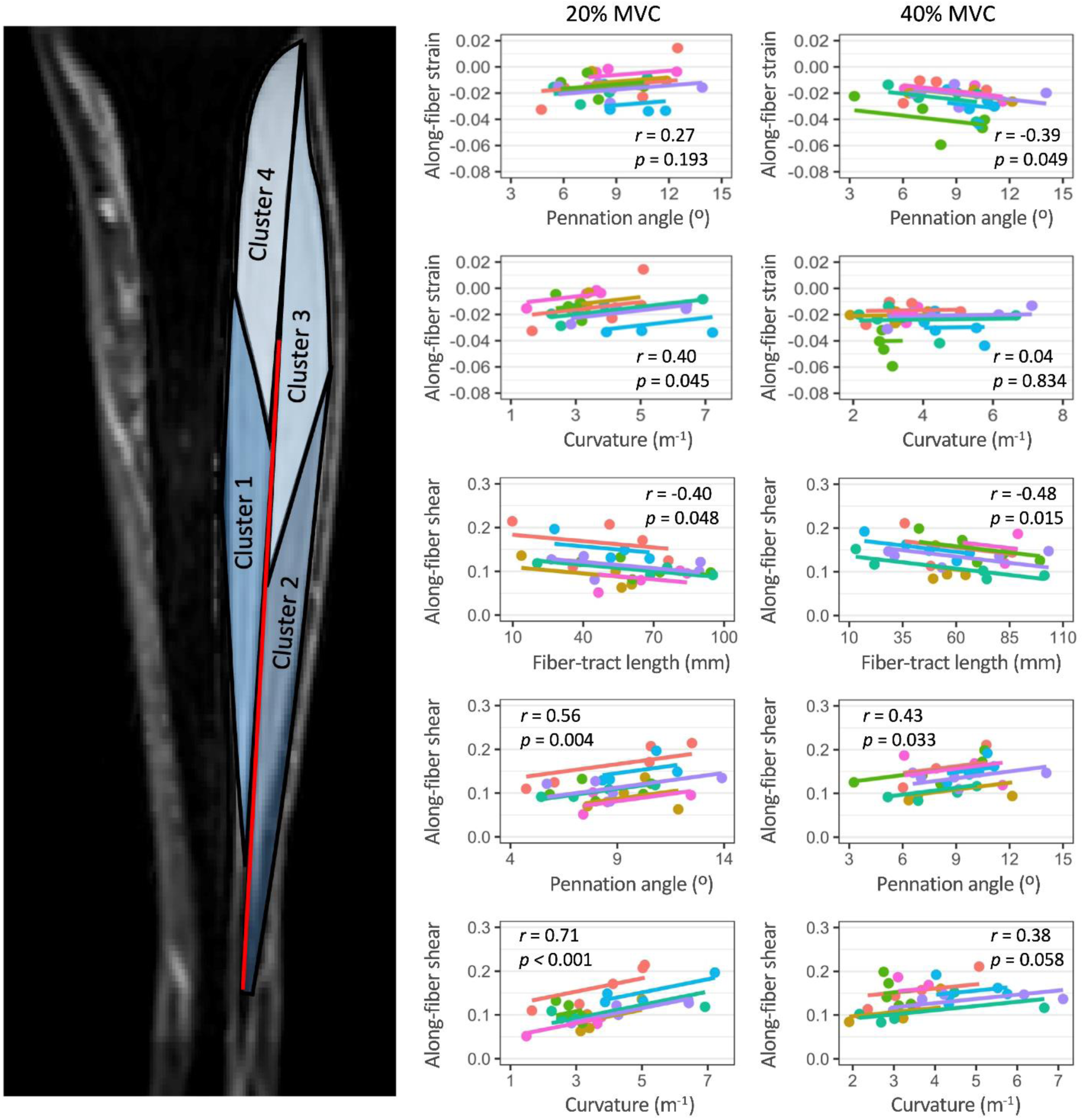
Repeated measures correlation plots showing the 20% and 40% MVC data of the architecture parameters and strain metrics that are significantly correlated. The TA muscle from each participant was divided into different fiber-tract clusters that included fiber-tracts with similar architecture, with each color representing a different participant and each point representing their respective fiber-tract clusters. The correlation coefficient, *r*, and the *p*-value of each correlation are shown in each plot. Notice how some variables are significantly correlated (*p*<0.05) at 20% MVC but not at 40% MVC and vice versa. The image on the left shows a longitudinal view of one TA muscle and how it was divided into different fiber-tract clusters.

**Table 5.** Correlation coefficients, *r* [95% confidence interval (CI)], and *p*-values of the repeated measures correlations between the mean strain metrics and the muscle architecture parameters of the fiber-tract clusters.

| Variables correlated |  | 20% MVC |  | 40% MVC |  |
| --- | --- | --- | --- | --- | --- |
| | | $r$ [95% CI] | $p$-value | $r$ [95% CI] | $p$-value |
| Along-fiber strain | $L_{FT}$ | -0.10 [-0.47 0.31] | 0.649 | 0.36 [-0.04 0.66] | 0.078 |
| | $\alpha$ | 0.27 [-0.14 0.60] | 0.193 | -0.39 [-0.69 0.00] | **0.049** |
| | $\kappa$ | 0.40 [0.01 0.69] | **0.045** | 0.04 [-0.36 0.43] | 0.834 |
| Along-fiber shear | $L_{FT}$ | -0.40 [-0.69 0.00] | **0.048** | -0.48 [-0.74 -0.11] | **0.015** |
| | $\alpha$ | 0.56 [0.21 0.78] | **0.004** | 0.43 [0.04 0.70] | **0.033** |
| | $\kappa$ | 0.71 [0.44 0.86] | **<0.001** | 0.38 [-0.01 0.68] | 0.058 |
| Cross-fiber shear | $L_{FT}$ | -0.32 [-0.64 0.08] | 0.116 | -0.01 [-0.40 0.39] | 0.966 |
| | $\alpha$ | -0.02 [-0.41 0.38] | 0.923 | -0.27 [-0.60 0.14] | 0.199 |
| | $\kappa$ | -0.08 [-0.46 0.32] | 0.690 | 0.06 [-0.34 0.45] | 0.768 |

## 4. Discussion

We present the first measurements of both 3D architecture and strain in a whole human muscle during moderate intensity (20-40% MVC) voluntary isometric contractions. These measurements are derived from the registration of structural images that show muscle bulging in the anterior direction during contraction (Figure 1), consistent with ultrasound-based observations (Raiteri et al., 2016). The decrease in fiber-tract length and increase in pennation angle found in the TA muscle during contraction agree with previous results from ultrasound in a limited field of view (Ito et al., 1998; Raiteri et al., 2016). Fiber-tract shortening reflects muscle fascicle shortening and is associated with the shortening of sarcomeres during contraction, while changes in pennation angle result indirectly from tendon lengthening. Moreover, we found that curvature increased in the TA during contractions, potentially resulting in increased intramuscular pressure (Otten, 1988; Van Leeuwen & Spoor, 1993). These findings are consistent with our first hypothesis and show the potential of this registration-based method to measure whole-muscle architecture during contraction.

The TA’s measured strain metrics are spatially heterogeneous during contraction. Intramuscular heterogeneity in strain rates has also been observed during contraction (Hooijmans et al., 2024; Malis et al., 2020). In the present study, larger intramuscular heterogeneity in along-fiber strain (*ε*) occurred at 40% MVC than at 20% MVC (Figure 3), demonstrating changes in intramuscular strain distributions with increased contraction intensity. The longitudinal heterogeneity in *ε* could be related to motor end-plate location (Drost et al., 2003), spatial heterogeneity in sarcomere shortening (Moo & Herzog, 2018), and other functional aspects of muscle. The effect of contraction intensity on the intramuscular heterogeneity of *ε* showcases how these aspects could vary at different contraction intensities. The results presented herein highlight the potential of a registration-based approach to investigate functional aspects of muscle activation and contraction more extensively.

Along-fiber shear (*B*_1_) was lower and cross-fiber shear (*B*_2_) was similar to predictions from computational models of the biceps brachii (Blemker et al., 2005). Low *B*_1_, in comparison to (Blemker et al., 2005), could be caused by the small changes observed in pennation angle during contraction. *B*_1_is thought to be related to force transmission from intramuscular connective tissue (Purslow, 2002), and could be used in future studies to investigate force transmission during different contraction intensities and types of contractions. The regional heterogeneity observed in *B*_2_ indicates that changes in shape perpendicular to the muscle fascicles are lower in areas of the TA superior to the aponeurosis (*No-Aponeurosis*). The interaction between the lengthening aponeurosis and shortening muscle fibers (Ito et al., 1998) potentially results in the higher *B*_2_ measured in the *Deep* and *Superficial* regions. Moreover, the highest *B*_2_ measured in the *Deep* region could be caused by the region’s interaction with the tibia, which is stiffer than the muscle and therefore constrains deformation in the axial plane. These results show the variation in strain along and within different regions of muscle during isometric contractions, consistent with our second hypothesis.

The region-specific changes in architecture and strain within the TA muscle underscore the importance of considering the heterogeneous architecture of skeletal muscle when conducting mechanical analyses. Computational simulations have shown that variations in muscle architecture influence strain nonuniformity (Blemker et al., 2005). To evaluate architectural variation within the muscles, we considered fiber-tract clusters with architecturally similar fiber-tracts where data could be averaged. The correlations between fiber-tract cluster architecture parameters and strain metrics show that pennation angle and curvature influence along-fiber strain at 20% and 40% MVC, respectively, similarly to previous reports (Azizi & Deslauriers, 2014). Moreover, the correlations between along-fiber shear and the architecture parameters (Figure 5) suggest that architecture could influence force transmission via the intramuscular connective tissue. Differences in the correlations between the same variables at 20% and 40% MVC could be attributed to the non-linear mechanical behavior of muscle’s extracellular matrix and warrant further study.

This study has several limitations, as image registration has yet to be validated against known measures of strain. Nevertheless, it demonstrated robustness at identifying heterogeneous deformation patterns in muscle (Karakuzu et al., 2023) and transforming muscle fiber-tracts under passive deformation (Hooijmans et al., 2025). Additionally, the methodology presented herein is limited to healthy participants and needs to be tested in pathological conditions, which present challenges in DTI data acquisition (Hooijmans et al., 2015), but could take advantage of high image contrast, i.e. fat-water contrast (Hooijmans et al., 2017), to ensure accurate registration. Finally, our registration-based methodology is constrained to static assessments and cannot quantify the viscoelastic aspects of contraction development over time.

## 5. Conclusions

We have presented the first estimations of both 3D architecture and strain in a whole human muscle during moderate (20-40% MVC) voluntary contractions. Our results highlight the heterogeneous changes in muscle architecture and strain in the TA during isometric contraction, and how muscle architecture relates to intramuscular variations in strain. Our results showcase the potential of MRI-based methods to obtain 3D estimates of whole-muscle architecture and strain during contraction, and the breadth of data provided by our methodology allows for new avenues of skeletal muscle research. This methodology can provide experimental validation to computational simulations of muscle contractions and be used to study the effect of neuromuscular conditions, such as muscular dystrophy, in whole-muscle architecture and mechanics.

## 6. Acknowledgements

The authors acknowledge support from NIH/NIAMS R01 AR073831 and the Stephens Family Clinical Research Institute.

