## Supplementary material for "MRI-based 3D Estimation of Skeletal Muscle Architecture and Strain during Contraction"

**S1. Schematics showing the regional subdivisions used to study the tibialis anterior (TA) muscle**

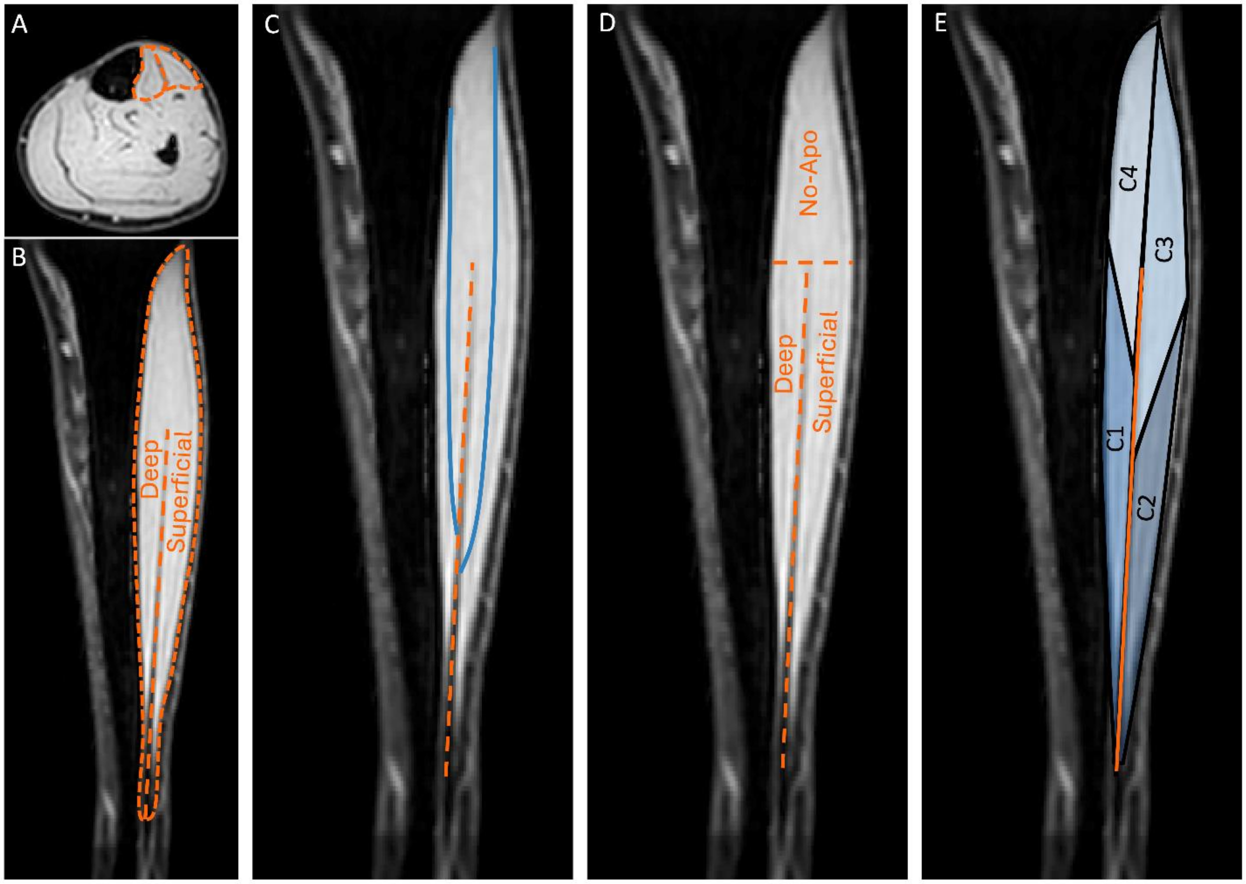


**Supplementary figure 1.** A) Axial anatomical image showing the outer boundary of the tibialis anterior (TA) muscle and a middle line passing through the aponeurosis, dividing the muscle into the *Deep* and *Superficial* compartments. B) Longitudinal view of the lower leg showing the outer boundary of the TA muscle and the aponeurosis dividing the muscle into the *Deep* and *Superficial* compartments. C) Longitudinal view showing the fiber-tracts generated from the aponeurosis (blue lines) into the *Deep* (left) and *Superficial* (right) compartments. D) Regional division of the TA muscle into regions corresponding to the *Deep* and *Superficial* compartments and the *No-aponeurosis* region located superior to the muscle aponeurosis. E) Division of the TA muscle into regions corresponding to different fiber-tract clusters that include fiber-tracts with similar architecture. Note how all regions from the fiber-tract clusters originate at the aponeurosis, where the fiber-tracts are seeded. The fiber-tract clusters were defined differently for each muscle, depending on the architectural similarity of the fiber-tracts within the muscle.

**S2. Creation of fiber bundles from fiber-tracts with similar architecture**

The fiber bundles were created using the algorithm previously described by (Damon et al., 2002), as adapted to account for the use of non-planar fiber-tract seeding methods. The fiber similarity threshold used was 0.5, with 20 neighboring fibers compared for fiber similarity, a maximum number of 130 fibers exceeding the fiber similarity threshold in each bundle, a minimum bundle size of 20 fibers and a maximum distance of 30 millimeters between the centroids of fibers in the same bundle.

### S3. Determination of image registration parameters for estimation of muscle deformation during contraction

Fiber-tract transformation and strain mapping require displacement fields that guarantee topology preservation. Water and out-of-phase (TE = 1.18 ms) images from the SPGR acquisitions during rest and contraction were registered at different levels of displacement field regularization by varying the *AccumulatedFieldSmoothing* (AFS) parameter in the *imregdemons()* function from 0.5 to 1.5 in step increments of 0.25. The effect of displacement field regularization and image-contrast were evaluated to find the best set of image registration parameters. The following image registration quality parameters were used to perform the evaluation:

1. Jacobian (J) of the muscle: defined as the determinant of the deformation gradient of the displacement field ($\vec{u}$), to evaluate the topology preservation (one-to-one mapping, $J > 0$) of the registration

$$J = \det\left(\frac{\partial \vec{u}}{\partial \vec{x}} + \boldsymbol{I}\right),$$

   where $\boldsymbol{I}$ is the 2$^{nd}$ order identity tensor

2. Dice-Sorensen Similarity Coefficient (DSC): defined as a measure of the overlap between the registered configuration of an object (A) and its target deformed configuration (B). Ranges from 0 (no overlap) to 1 (complete overlap). Evaluated in the registered muscle and aponeurosis masks and their respective manually segmented pair obtained from the contraction images

$$DSC = \frac{|A \cap B|}{|A| + |B|}$$

3. Euclidean distance: defined as the mean Euclidean distance between corresponding points of the registered aponeurosis mesh and the contracted aponeurosis mesh
4. Modified Hausdorff distance: defined as an indirect measure of the overlap between the registered and contracted aponeurosis mesh, based on the distance between the points of the registered and contracted aponeurosis mesh (Dubuisson & Jain, 1994)

J was evaluated for the different values of AFS in both the water and out-of-phase images. The minimum value of J was computed in each muscle to evaluate the topology preservation of the registration's displacement field $\vec{u}$. We found that topology was not preserved in the registration of the 40% MVC images in at least one of the participants at an AFS of 0.5, 0.75, and 1. The minimum J computed in each muscle gradually increased with increasing AFS, resulting in diffeomorphic mappings from the registration of the 20% and 40% MVC images from all participants at an AFS of 1.25 and 1.5. The mean J ranged between 0.97 and 1 in the displacement fields obtained from the mappings of the contractions at 20% and 40% MVC. These results demonstrate volume preservation of the muscle during isometric contraction, as expected. Additionally, the maximum J computed in each muscle decreased as AFS increased, because of increased displacement field regularization. These results were observed at both water and out-of-phase images (Supplementary figure 2). We decided to use an AFS of 1.5 in this study, as it guaranteed topology preservation and had more displacement regularization than an AFS of 1.25. A previous study has found that strain map computations are robust to image registration parameters (Karakuzu et al., 2023), supporting that our decision to use an AFS of 1.5 would not bias our results.

To decide between the water and out-of-phase images, we compared the DSC of the muscle and aponeurosis masks, the Euclidean distance, and the Modified Hausdorff distance from the registration of the two different image contrasts. The median muscle DSC was within ±0.5% of 0.94 for out-of-phase and water images at 20% and 40% MVC. Median aponeurosis DSC was 0.848 for out-of-phase images and 0.844 for water images at 20% MVC, a 0.5% difference. At 40% MVC, there was a 1.5% difference between the median aponeurosis DSC of 0.801 for out-of-phase images and 0.813 of water images. The median Euclidean distance was 3.35 mm in the out-of-phase images and 3.26 mm in the water images at 20% MVC, while it was 3.48 mm in out-of-phase images and 3.42 mm in water images at 40% MVC. The median modified Hausdorff distance was 1.99 mm in the out-of-phase images and 1.93 mm in the water images at 20% MVC. At 40% MVC, the median modified Hausdorff distance was 2.06 mm in out-of-phase images and 1.70 mm in water images. Even though the muscle and aponeurosis DSC were within a 2% difference between water and out-of-phase images at 20% and 40% MVC , the median Euclidean distance and modified Hausdorff distance were lower in the water images images (Supplementary figure 2). Thus, we decided to use water images for our muscle architecture transformation and strain mapping analyses. The image registration quality parameters obtained from isometric contraction represent an improvement with respect to the parameters obtained from passive deformation (Hooijmans et al., 2025) and are on par with those found in a previous study that used MRI-based image registration to measure strain in intervertebral discs (Yoder et al., 2014).

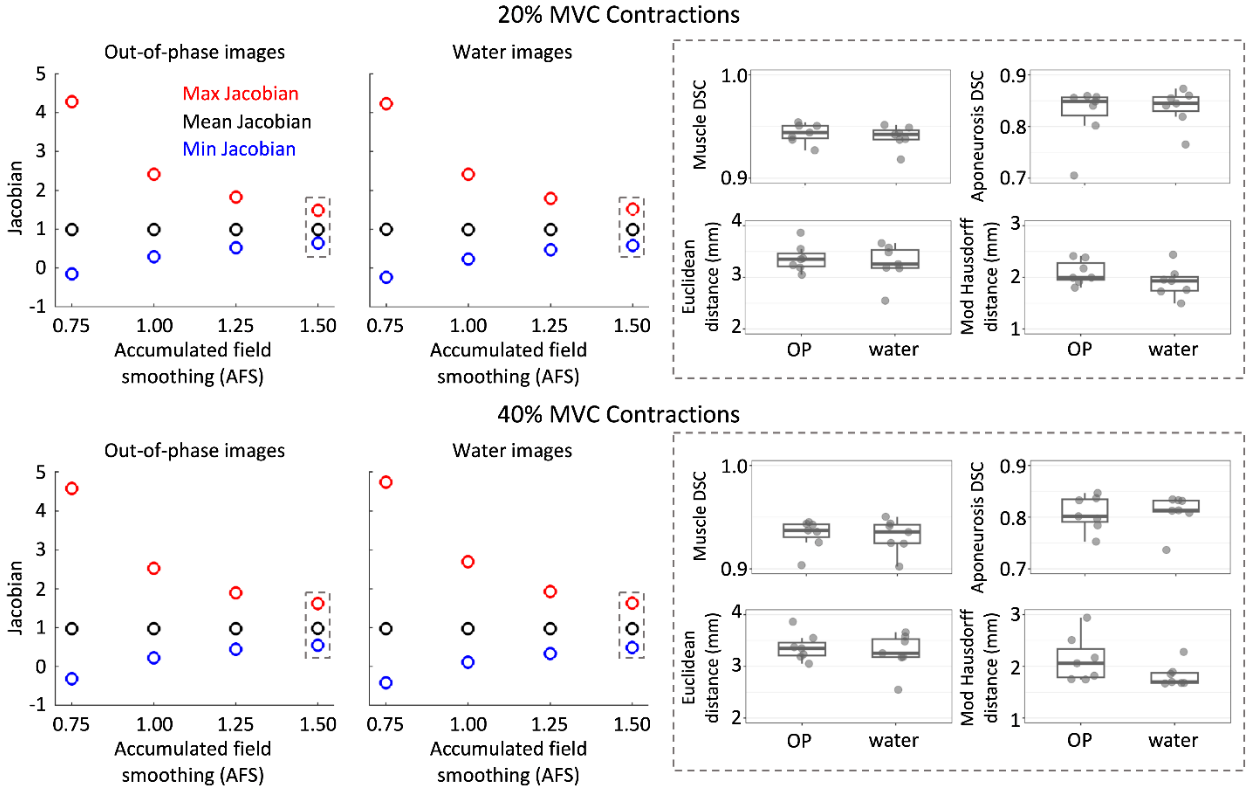


**Supplementary figure 2. Estimates of the image registration quality parameters used to optimize the image registration of the rest and contracted muscle images.** The Jacobian plots on the right show the median measurements of the maximum, mean, and minimum Jacobian obtained from the registration of the out-of-phase (OP) and water images of the seven participants at 20% and 40% MVC for AFS of 0.75 to 1.50. The results of an AFS of 0.5 are not shown because the maximum Jacobian was higher than 5 and the minimum Jacobian was lower than -1 for all participants. The box and dot plots on the left show the muscle and aponeurosis DSC, Euclidean distance, and Modified Hausdorff distance measured from the registration of the images of the seven participants.
